# Heterospecific aggression differs predominately by species, rather than sex, in Lake Malawi cichlid fish

**DOI:** 10.1101/720375

**Authors:** Emily C. Moore, Reade B. Roberts

## Abstract

Because of their striking diversity, Lake Malawi cichlid fish have been well studied for male aggression, particularly among dominant males of closely related sister species within the framework of mate-choice and speciation. However, aggression in females has been largely ignored, and variation in aggressive behaviors between more distantly-related taxa is not well understood despite its potential impact in a complex community structure. To better understand variation in patterns of aggression between species, we presented males and females from five species of Lake Malawi cichlid with a non-predator intruder and recorded all movement and aggressive acts. Additionally, we measured excreted cortisol levels the day after the intruder assay to evaluate one physiological aspect of stress response. We identified species-specific patterns in both specific aggressive acts, and overall level of aggression. Additionally, we found that sexual dimorphism in aggressive acts varies by species and act, where the species with the most aggressive males also has aggressive females. Additionally, cortisol levels vary by taxa, and are associated with restless behavior in the intruder assay, but not levels of aggression. These findings have bearing on understanding sex differences in aggression and their impact on community structure in this important model of rapid evolution.

## Introduction

In many animal species, aggressive behaviors are used to resolve competition over access to resources that influence fitness, such as mates, food and space^1,2^. While the focus of most aggression studies is on aggression between members of the same species, heterospecific aggression is common across animal taxa, and species can vary in their relative levels of conspecific and heterospecific aggression^3^. In complex, species-rich communities, heterospecific aggression depends on environmental factors such as predation^4^, chemical signals from conspecifics^4,5^, habitat type^6^, and overlap in dietary specialization^7^.

African cichlid fishes are an excellent evolutionary model for exploring species-level diversity in heterospecific aggression; Lake Malawi has regions with rich communities where many, recently diverged species coexist^8,9^, and food and space resources are finely partitioned into definable microhabitats^8,10^. In these communities, there are sex differences in territoriality—-dominant males hold stable territories which are used for feeding and attracting mates, while subordinate males and females inhabit a wider range surrounding these territories^10,11^. Territory quality can influence reproductive success in cichlids^12^, making successful territory defense important for both feeding and successful breeding. Given community structures where males of different species have territories very close to one another, the combination of high levels of male aggression towards conspecifics and low levels of aggression towards heterospecifics has been theorized to play an important role in the evolution and maintenance of diverse, closely-related sympatric species during speciation^13^.

However, the presence of sympatric sister species does not necessarily result in a reduction in heterospecific aggression; wild populations of brook stickleback (*Culaea inconstans*) in communities with sympatric ninespine stickleback (*Pungitius pungitius*) have been shown to be more aggressive than those from populations where *Culaea* were the only stickleback species present^14^. Even in Lake Malawi, where low heterospecific aggression is an important facet of the proposed mechanism for sympatric evolution^13^, there is evidence that sister species pairs differ in levels of heterospecific aggression, and this variation seems to be associated with adaptation to different substrate type, or microhabitat^6^. Given that microhabitat is the same between sexes, this may indicate that ecological selection, rather than sexual selection, predominately shapes heterospecific aggression in this group.

In this paper, we describe and quantify heterospecific aggression in both sexes of five Lake Malawi cichlid species, in order to evaluate whether heterospecific aggression may be decoupled from sex. We also identify species differences in pattern and amount of aggressive behavior, as well as a physiological measure of stress, the hormone cortisol. These five species represent fine-scale variation between sister species within the rock genus *Metriaclima* [*M. aurora, M. lombardoi, M. callainos*, and *M. pyrsonotos*], as well as the more distantly related, sand-dwelling species *Aulonocara baenschi*.

## Results

### Aggression differences by sex and species

At the broadest levels, behavioral patterns can be visualized with a time-line representation of the ethogram, or catalog of all the behaviors observed in the assay (figure S1), which allows for a non-quantitative overview of the patterns of behavior displayed by each individual fish. A comparison of mean summary values for each behavior (count or duration) by species, sex, and the species by sex interaction showed differences in the most aggressive behaviors. The aggressive behaviors lunging (F(9,61)=3.71, corrected p=0.0063) and chasing (F(9,61)=2.84, corrected p=0.044) have significant species main effects, as did the attentive fin display behavior (F(9,61)=4.69, corrected p=0.0008). Sex and the interaction term were left in the model despite non-significance because *M. lombardoi* shows aggressive behaviors in males but not females; while this may be an artifact, the pattern is clear for three behaviors, and thus deemed biologically relevant and may be detectable with a greater sample size (figure 1, top three panels). The quiver display was not significantly different by species in the model (F(9,61)=2.36, corrected p=0.117), but was notably only present in 4 of 8 *M. aurora* males, and a single *M. lombardoi* male, and thus the only male exclusive behavior recorded (figure 1, third panel from the top).

**Figure 1.**
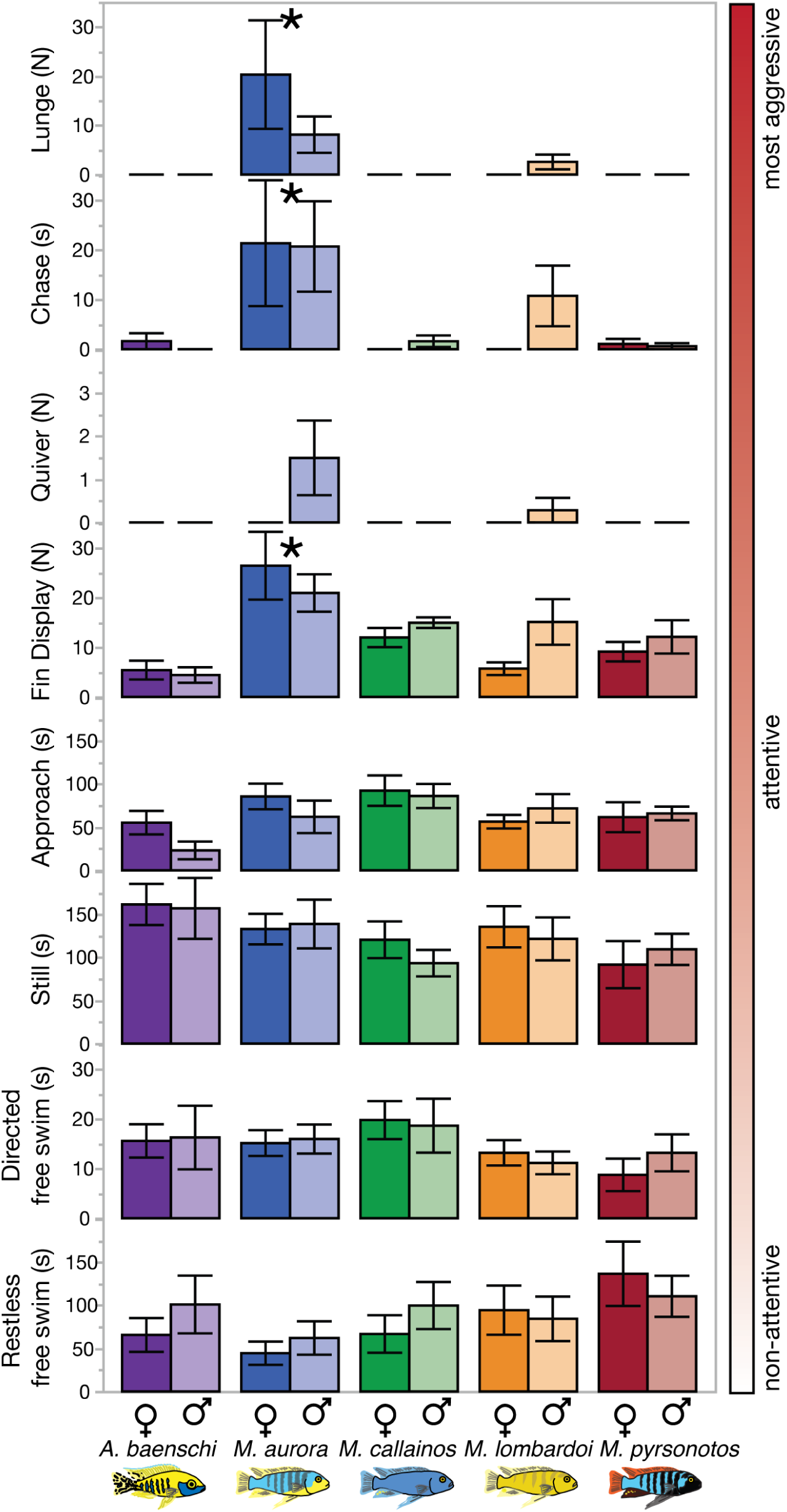
Movement and action states by sex and species. Duration of movement activities (s) and counts of actions (N) of focal fish relative to a *Labeotropheus trewavasae* intruder for both sexes of five species (n=7 per sex). Activities are organized from non-attentive to most aggressive. Note that the majority of behaviors were seen in both sexes, with the exception of the quiver display (present only in *Metriaclima aurora* and *M. lombardoi* males).

The most aggressive species, *M. aurora*, and the least aggressive species, *A. baenschi* display markedly different mean values for high-aggression behaviors (see top four panels, figure 1), but remarkably similar values for relatively low aggression behaviors (see bottom four panels, figure 1). This supports that the differences in aggression are not merely due to differences in overall activity, but species-specific responses to a *Labeotropheus* intruder.

There were no species differences in the amount of time spent focused on the intruder (attentive and aggressive motions) relative to the total time; on average, fish spent almost twice as much time focused on the intruder (197.73±7.8 s) as they spent doing non-attentive motions (102.03±7.9 s). For both of the non-attentive movement behaviors, there were no detectable species or sex differences in time spent free swimming (figure 1, bottom three panels). All species spent less time engaged in directed free swimming (14.77±1.2 s) than restless free swimming (87.26±8.3 s).

Intruders did not display any of the attentive or aggressive acts, including approach and chase. The proportion of time the *Labeotropheus* spent in the territory was lower than all resident species, and did not differ by the species of the resident (supplemental figure 2B).

### Temporal and spatial differences in aggressive acts

Some of the behaviors changed in frequency over time, where resident fish spent more time swimming restlessly (figure S2A, top panel), and less time in the territory (figure S2B, top panel) as the assay progressed. There were species differences in resident territory occupancy, with *A. baenschi* spending the most time in or very near their territories (figure S2B, top panel). The intruders spent less time in the territory overall, and were found in the territory region in a time-independent, species-independent manner, providing a control value for territory time expected for a non-habituated fish (figure S2B, bottom panel). Additionally, the momentary aggression score showed an increase over time for *M. aurora*, where differences are not detectable in the first minute, but became pronounced after the second (figure S2C).

The knowledge of position and action simultaneously allowed for comparisons of whether behaviors were more common inside or outside of the territory. No clear species patterns were found for the most aggressive acts; however, the attentive fin-spread display showed an intriguing spatial difference by genus. *Aulonocara baenschi* was much more likely to perform the fin-spread display while inside the territory, and the *Metriaclima* were more likely to perform the display while swimming around the open regions of the tank (figure 2). This may suggest that fin-spreads were defensive for *A. baenschi* and offensive for *Metriaclima*, even for those *Metriaclima* with low overall aggression, including *M. callainos* and *M. pyrsonotos*.

**Figure 2.**
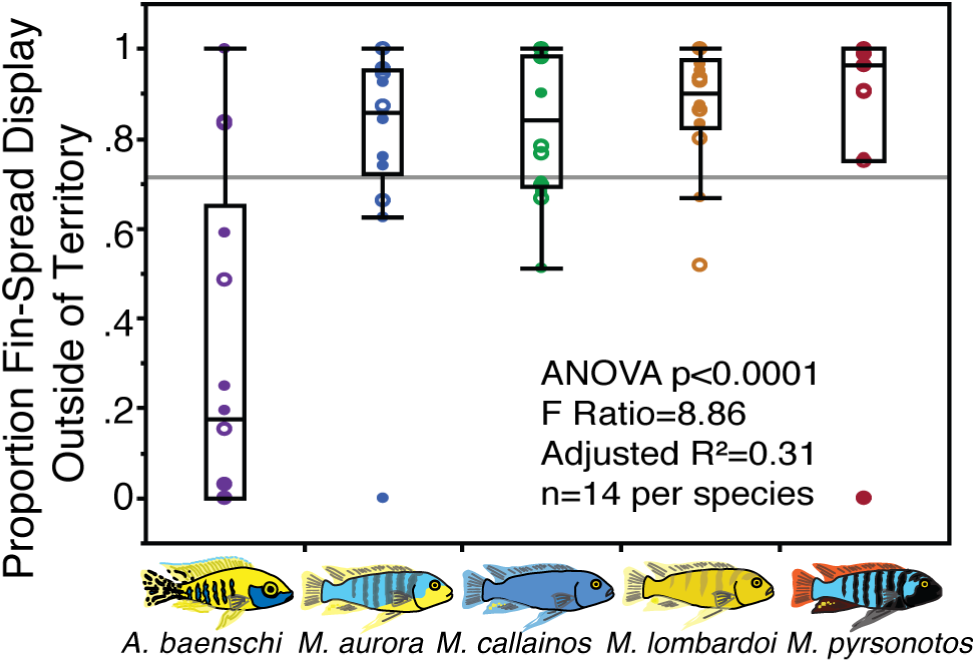
Relative territory position of fin-spread display behavior. Proportion of fin-spread displays performed outside of the territory region of the aquarium. A value close to 1 indicates that displays were made away from the flower pot territory, and a value closer to 0 indicates that displays were made in the flower pot territory. All four *Metriaclima* species predominantly displayed from outside the territory, while *A. baenschi* displayed from inside the territory. This metric does not take number of displays into account, only position in the tank while the display was made. Male data points are represented by closed circles, female data points are represented by open circles.

### Transition patterns between behaviors

A comparison of mean transition probability diagrams between the most aggressive species, *M. aurora* (figure 3A, top), and the least aggressive species, *A. baenschi* (figure 3A, bottom), reveals some conspicuous presence/absence differences in transitions that can allow for insight into how these species responded to intruders in their space. For instance, when *A. baenschi* were restlessly free swimming, they never initiated chases, and conversely never returned to restlessness directly after chasing, as did the *M. aurora*. Instead, *A. baenschi* transitioned through an approach motion before chasing, indicating that they required a build-up of attentiveness before the aggressive act of chasing intruders. Some transitions, however, differed very little. Once either species approached the intruder, it had a high probability of becoming still, rather than chasing the intruder.

**Figure 3.**
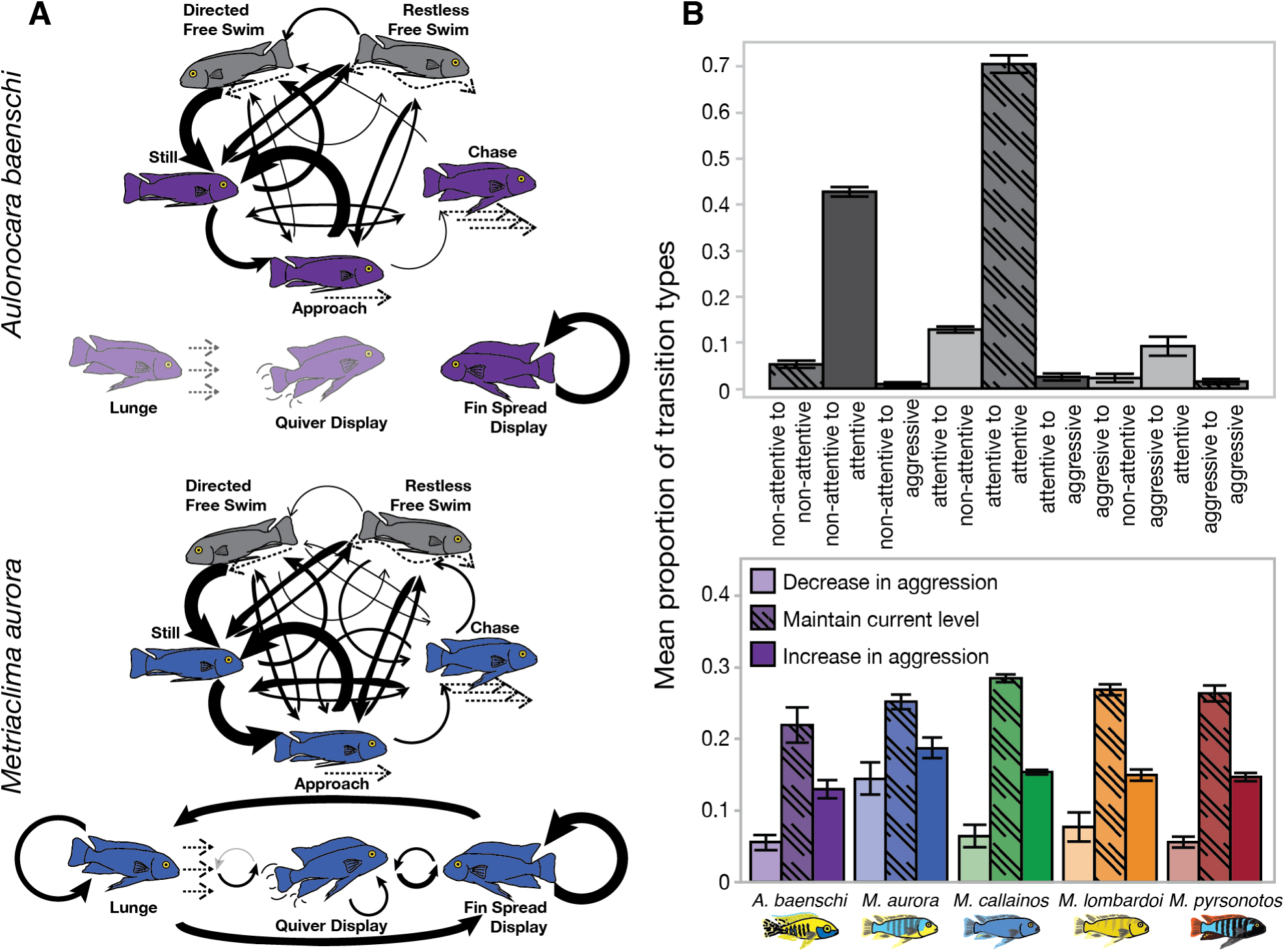
Behavioral transitions by activity and aggression level change. Average transition probability by activity state for *A. baenschi* (A, top panel) and *M. aurora* (A, bottom panel), n=14 per species. Movement and action behaviors are unconnected by arrows because they are of different behavior classes, and occurred independently of each other. Greyed out behaviors indicate that lunge and quiver display were not performed by *A. baenschi*. Relative bolding of arrows indicates the probability that, once in a behavioral state, the fish will move to the other state The transitions can be categorized by type of transition (B, top panel, all fish from all species included), or whether the change decreases, increases, or maintains current aggression levels (B, bottom panel). For all species except the most aggressive (*M. aurora*), decrease in aggression level is the least common change, and maintenance of aggression level is the most common.

Notably, if a fish was in a non-attentive state it was most likely to transition to an attentive state, and once in an attentive state, it was most likely to stay attentive (figure 3B, top). Overall, transitions to aggressive behaviors were relatively rare. When the transitions are categorized by relative increase or decrease in aggression state, *M. aurora* has a slightly different pattern than the other four species. *Metriaclima aurora* were just as likely to increase aggression as decrease it, reflecting a pattern of transitions where increased switches from attentive to aggressive activities were common (figure 3B, bottom).

### Cortisol level differences by species

We measured excreted cortisol levels to serve as a physiological measure of basal stress response, taken the day after the resident-intruder assay. Excreted cortisol levels differed at the genus level, with *A. baenschi* possessing the lowest levels (figure 4A). Despite a prediction that aggressive individuals would have higher cortisol levels, we found no association between aggressive acts and cortisol. However, the stress hormone was correlated with other attentive and non-attentive behaviors; individuals with higher excreted cortisol levels had an increased number of instances of approaching the intruder (4B), swimming restlessly (4C), and staying still (4D). The increased number of instances for these three behaviors indicates that fish with higher cortisol switched from restless swimming to attentive acts more often. While it is possible that these fish were simply more active in general, they were not more likely to have an increased number of instances swimming toward their territory (4E). All together, we interpret these data as indicating a connection between physiological stress and increased transitions between attentive and restless behavior.

**Figure 4.**
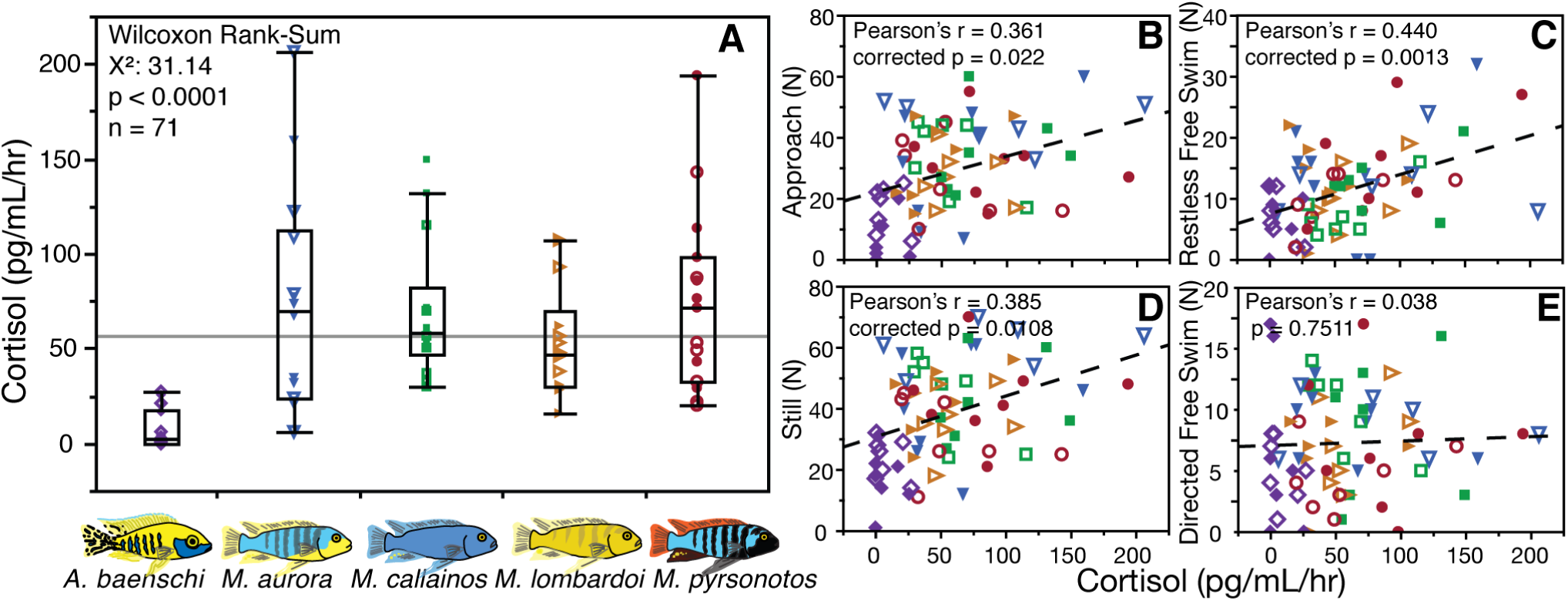
Species differences in excreted cortisol levels, and corresponding association with non-attentive restless swimming behavior. Excreted cortisol levels are significantly lower in *A. baenschi* than in *Metriaclima* (A). The level of excreted cortisol is correlated with the instances of approach in the intruder (B), instances of restlessly swimming (C), and instances of being still (D), but not the number of times a fish swims in a directed manner (E). Male data points are represented by closed shapes, female data points are represented by open shapes for reference, though no sex differences in cortisol were detected. Shapes indicate species: *A. baenschi*, diamond; *M. aurora*, down-pointed triangle; *M. callainos*, square; *M. lombardoi*, right-pointed triangle; *M. pyrsonotos*, circle.

## Discussion

Much of the research about animal heterospecific aggression has found higher levels of aggression between more closely related taxa. A meta-analysis of 126 studies of heterospecific aggression showed that, controlling for levels of conspecific aggression, confrontations between species of the same genus had higher heterospecific aggression levels than those of different genera^3^. Closely related species may give similar signals, including visual (i.e. color, or shape) or chemosensory forms of species recognition. Dijkstra and Groothuis surveyed studies of male aggression in cichlids, and found that males were more likely to be aggressive to other males of the same color, stabilizing diversity in male color and allowing for sympatric coexistence of many (differently-colored) species^13^. Despite this overall pattern, the authors note that a small number of species tested show biases against different-colored males and acknowledge that aggression between sister taxa can be asymmetric. However, this explanation is based predominately on studies that used comparisons between males of sister taxa that differed in nuptial coloration to examine interactions of species in sympatry, and so are missing broader comparisons of aggression between more distantly related species. Indeed, Pauers and colleagues compared two differently pigmented sister species and found that the focal species, *Metriaclima mbenjii*, performed more lateral displays to conspecifics than the heterospecific *M. zebra*, as put forth by the model proposed by Seehausen and Schluter^15^, but also showed a trend towards performing more aggressive bites towards the more distantly related Labeotropheus fuelleborni^16^. The finding that specific acts may vary depending on the species involved in the interaction is consistent with our findings, thus we suggest that when investigating heterospecific aggression, a nuanced examination of behavioral patterns should be used to capture relevant biological differences in how species are interacting.

Competition over similar resources can also potentially increase aggression between more closely related species^3^, where aggression is tuned towards heterospecifics that are adapted use to similar ecological resources. This has been shown for food resources in Lake Malawi, where cichlid species showed higher levels of aggression to heterospecifics when they had similar dietary specialization to the focal species, and lower levels of aggression to heterospecifics when they were dietary generalists^7^. In contrast, our study found variation in heterospecific aggression within the dietary generalist genus Metriaclima, indicating that ecological selection pressures on heterospecific aggression are likely more complex than partitioning of food resources alone. Since our fish were raised in a controlled, common garden setting, any differences in behavior we detect likely have a genetic basis, rather than being caused by behavioral plasticity in response to environmental conditions or microhabitat.

One major ecological factor that might explain behavioral variation in aggression between species is variation in microhabitat. Danley found that variation in substrate complexity, or microhabitat, predicted heterospecific, but not conspecific, aggression levels in the wild and in the lab^6^. Our study included one of the high heterospecific aggression species, *M. aurora* and one of the low heterospecific aggression species, *M. callainos*, examined by Danley and we replicated his findings; this provides further support to the idea that microhabitat use, rather than male-male competition or food resource partitioning, is driving variation in heterospecific aggression.

Cichlids are not the only species in which levels of heterospecific aggression are associated with divergence in microhabitat use. For another freshwater fish community, heterospecific aggression between stream fishes is related to overlap in microhabitat use during certain times of the year^17^. In the salamander species *Plethodon cinereus*, wild assortatively-mating color morphs have differences in male aggression, where striped males are more aggressive and more likely to hold leaf litter territories than non-striped males. The non-striped males’ non-territorial roaming behavior is associated with increased resistance to desiccation, and may be a case of niche partitioning resulting in speciation^18^. The association between aggression and space use could develop in a few ways, including territoriality/aggression mediating shifts in niche use due to exclusion of the less aggressive species^3^, or selection for increased levels of aggression of after niche partitioning occurs. We do not currently have the data to distinguish between these two possibilities, and this is an avenue for further research.

Perhaps given the prominence and visibility of male aggressive displays during courtship and territory guarding^1^, both heterospecific and conspecific female aggression is less often reported in the literature. When females are included in studies, they also show variation in aggression levels and types of aggressive acts. In wild-caught populations of the poeciliid fish *Brachyrhaphis episcopi*, females from low predation populations were more aggressive than males, though males performed more displays when shown a mirror^19^. Males and females of the cooperatively breeding Tanganyikan cichlid *Neolamprologus savoryi* both have high levels of heterospecific aggression against a predator species, but only have high levels of conspecific aggression during interactions with fish of the same sex^20^. South American convict cichlids (*Amatitlania nigrofasciata*) showed no sex differences in propensity to aggression after a startle, but did show sex differences in the types of aggressive acts^21^. Our findings support a model where female aggression is variable, types of behaviors can vary by sex (such as the quiver display), and, for 4 of 5 species, heterospecific aggression levels are the same in both sexes.

It is unclear whether the high occurrence of female aggression in studies where it has been examined indicates there is a biological trend for species that have aggressive males to also have similarly aggressive females, or if there is an ascertainment bias where researchers are more likely to include females in a study if they a priori know them to be aggressive. In the case of this study, we had no specific predictions regarding female aggression, other than an informal expectation that female aggression would be lower than male aggression since males hold territories in the wild, and females do not.

Variation in aggression at the individual level can be attenuated or increased by physical traits, such as relative size^22^, and social rank^23^. Though large size and social dominance are most often seen in male cichlids, experimental manipulation to create female-only communities of *Astatotilapia burtoni* in the lab results in females displaying male-like behaviors and hormone profiles^24^. Experimental evolutionary model systems allow for mechanistic dissection of behavioral traits such that we can begin to distinguish genetic modifiers of aggression from those mediated by social environment. Cichlids are one of these systems, where we can disentangle sex, social environment, and genetics/evolutionary history to understand how they contribute to variation in behaviors, including heterospecific aggression.

Beyond environmentally responsive cues that allow for individual flexibility in aggression due to social context, genetic modifiers of aggression that allow for stable inheritance of species-specific (potentially sexually dimorphic) behavioral patterns can either be autosomal, or at a sex locus (hormonal and genetic mediators of aggression reviewed in^25^). There is support for an autosomal genetic basis for aggression levels in teleost fishes decoupled from hormone levels; in sex-changing sand-perches (*Parapercis cylindrica*), aggression levels remain constant through the shift from female to male such that aggressive females become aggressive males, which indicates potential genetic constraints on sex-specific aggression^26^. Polygenic sex determination systems in cichlids^27–29^ may function in a similar way, allowing for flexibility in populations/species for aggression to be associated with sex, depending on whether the genetic modifier of aggression is linked to a sex locus that has effects that can be masked by the presence of one or more additional sex loci. This can functionally shift a once sex-specific trait into a trait present in both sexes, and provide a mechanism for both sexual and natural selection to act on a locus.

Mechanistically, hormones and other small signaling molecules have been associated with male aggression^25^. Naturally occurring, social role-associated variation in levels of aromatase (the enzyme that converts testosterone into estradiol) regulates aggression in the cichlid *Astatotilapia burtoni*^30^. *In the teleost fish Gymnotus omarorum*, aromatase can modulate non-breeding related aggression in a manner independent from the gonad, and in the face of low plasma hormone levels^31^, which may indicate that genetic modifications to this pathway might be able to influence aggression in both sexes. Signaling peptides also play a role in aggression in cichlids; additional studies using the *A. burtoni* model found no association of aggression with plasma levels of testosterone or 11-ketotestoterone, but found that a reduction in the regulatory peptide somatostatin increases aggressive acts in males^32,33^. Also, lower levels of the stress hormone cortisol have been associated with higher levels of aggression; less aggressive, non-territorial *A. burtoni* males had higher plasma cortisol levels^34^, and aggressive convict cichlids had lower post-assay excreted cortisol levels^35^. Both of the experiments evaluated conspecific aggression, so it is possible that cortisol is only associated with conspecific aggression or mediation of social dominance, rather than aggression per se.

The behavioral interactions of any community are complex, and influenced in different ways by both sexual and natural selection. Indeed, aggression in cichlids may play an important role in explaining the rapid diversification of sister taxa, particularly when natural variation in aggression towards others in the community is considered. The species-rich cichlid radiation can be a useful and informative system for contributing to the synthesis in the understanding of the evolution of social behaviors, both proximate (genetic and environmental modulation of neurochemicals and neural circuits) and ultimate (habitat, social structure) in an evolutionary framework^36^, especially if we include the great variety of behaviorally diverse species in the group. With these tools, we can harness naturally evolved differences in aggressive phenotype for understanding proximate mechanisms that result in behavioral diversification— not just overall levels of aggression, but how patterns of aggression are fine-tuned through adaptation.

## Methods

### Animals

Laboratory cichlid lines were wild-derived from animals collected in Lake Malawi, and maintained in according to the Institutional Animal Care and Use Committee (IACUC) guidelines in our aquaculture facility 1. Age matched individuals were raised in mixed-species groups in a 473-liter (184 cm x 47 cm x 60 cm) aquarium until onset of reproductive age at 6 months, when behavioral assays began. For each species tested, individuals from at least two families were placed into co-culture for testing. Following behavioral experiments and hormone collection, fish were anesthetized with buffered 100 mg/L tricaine methanesulfonate (MS-222) until cessation of opercular movement. During anesthesia, we recorded standard length (from the tip of the rostrum to the base of the tail, not including fin tissue) and weight, took a genital photograph for non-lethal sexing^37^, as well as a small sample of fin tissue for future DNA extraction, and injected a biopolymer tag to maintain identity in group housing conditions (Visible Implant Elastomer, Northwest Marine Technologies). Fourteen fish per species were assayed, (n=7 per sex per species, n=70 total).

### Housing Conditions

Individual animals were each placed in a 38-liter (51 cm × 28 cm × 33 cm) aquarium with a single flower pot territory for three days of acclimation to individual housing. Two banks of five aquaria were used to permit animals to maintain continuous visual access to other experimental fish housed singly in surrounding aquaria, but not to non-experimental fish housed in groups (see figure 5A). Because a maximum of 10 fish could be assayed at a time, a blocked design was used to ensure even distribution of species and sexes over each experimental day. Fish were not housed next to members of the same species. All fish were determined to be active and appetitive during the acclimation period, as judged by readiness to feed in the morning and evening prior to the assay. Following the intruder presentation, fish were maintained for an additional 48 hours in isolation territories, with a single 30-minute novel object assay involving exposure to a snail shell at 24 hours (data not included in this publication). Prior to returning individuals to group housing conditions, fish were placed in beakers for hormone collection.

**Figure 5.**
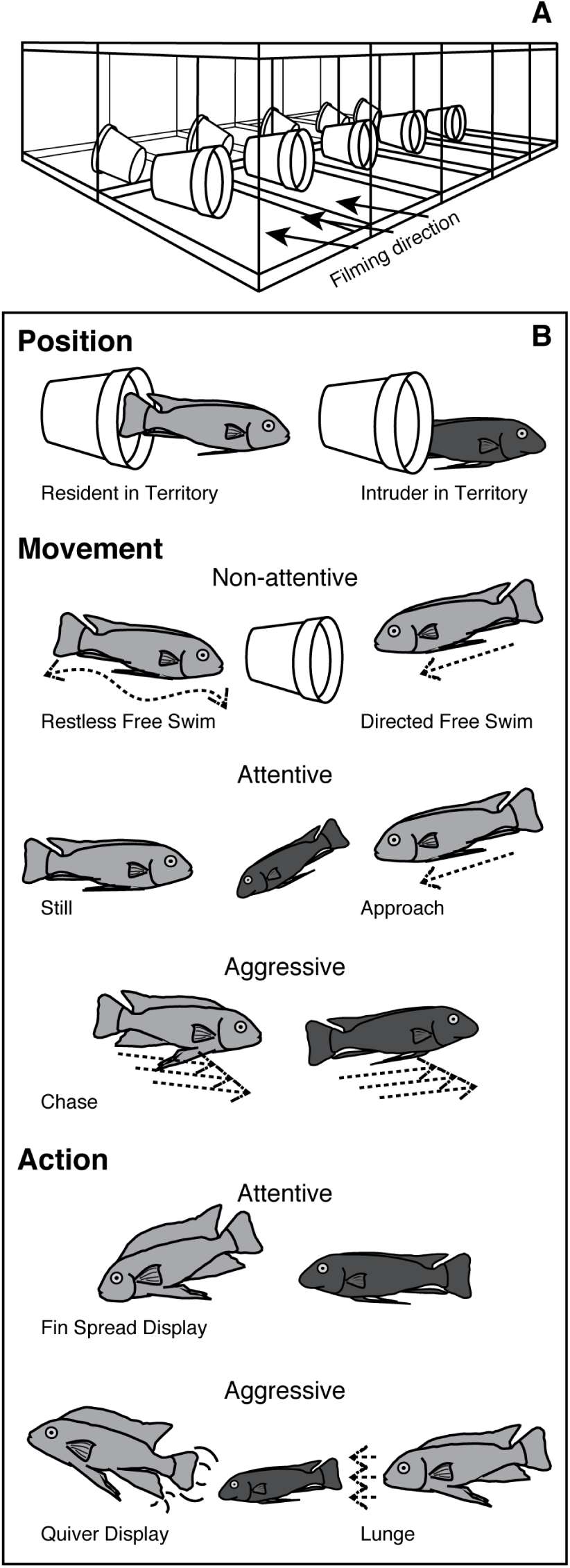
Experimental set up and recorded behaviors. Fish were individually housed in a bank of ten aquaria with visual access to their neighbors. Video was recorded for five minutes from the front aspect of the aquarium (A), and three separate aspects of behavior (position, movement, and action) were scored upon re-watching the videos (B). Position relative to territory was recorded for both the resident (focal) fish and the intruder fish. Mutually exclusive movement states were characterized as non-attentive, attentive, or aggressive based on the level of interaction with the intruder fish. Actions were characterized as attentive or aggressive, and could occur independently of movement state; for instance, a lunge is a short burst of activity, which could happen while the fish is still or swimming, and is distinct from a prolonged chase. Movement that was not towards the intruder was characterized as either restless free swim (where the focal fish swam back and forth aimlessly), or directed free swim (where the focal fish was headed to a specific place, usually the flower pot territory).

**Figure 6.**
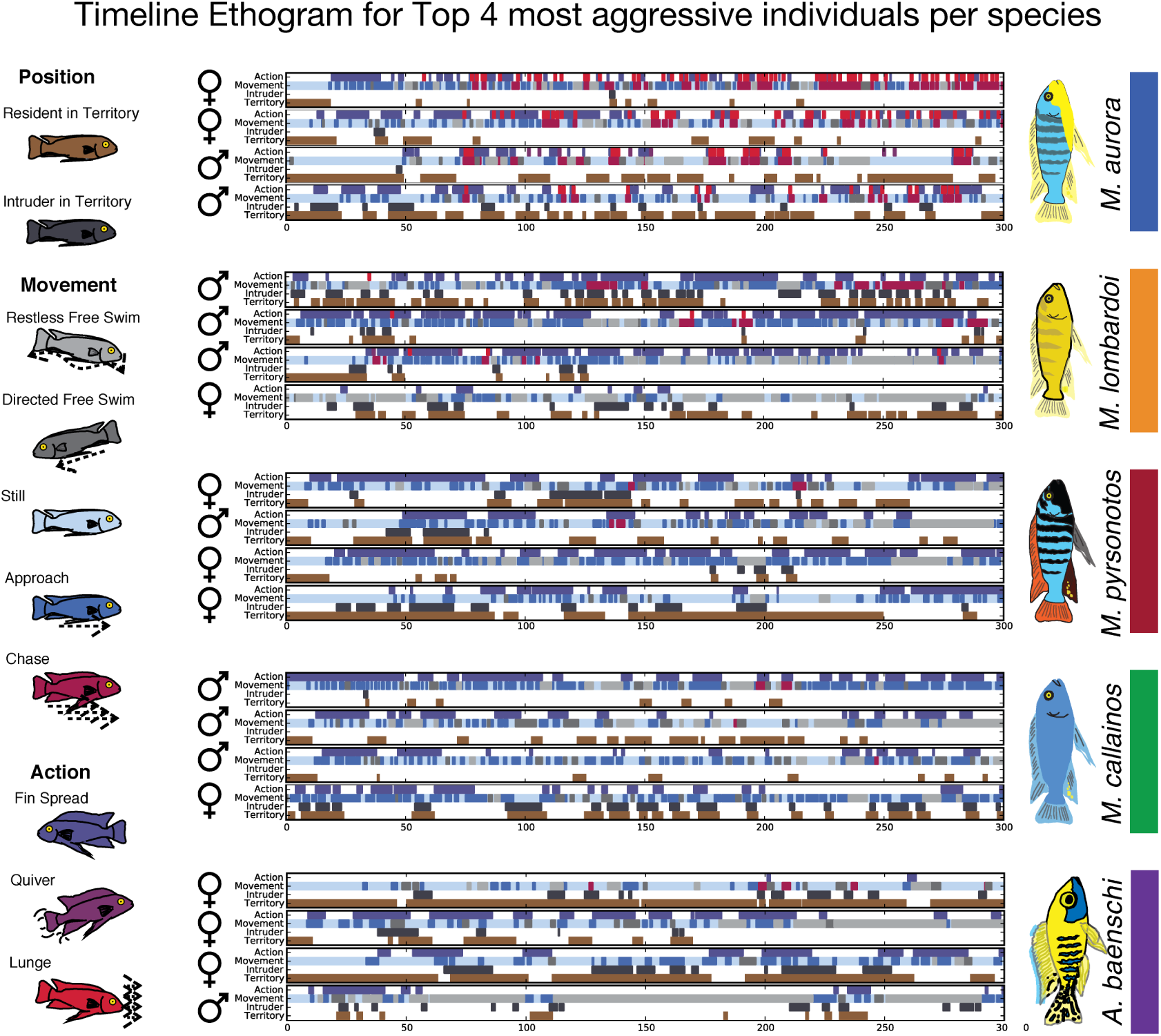
Figure S1. Ethogram time lines for the top four most active fish per species. Graphical overview of each of the top four most active fish per species for all measured behaviors over time; actions (top row) and movements (second row) are color-coded as indicated by the legend fish to the left, where gray indicates non-attentive behaviors, blues indicate attentive behaviors, and reds indicate aggressive behaviors. Position relative to the territory is noted for both the intruder (third row, gray) and focal fish (bottom row). The sex of each individual is included to the left of its time line; note that both males and females are included in the top four most active for all species tested.

**Figure 7.**
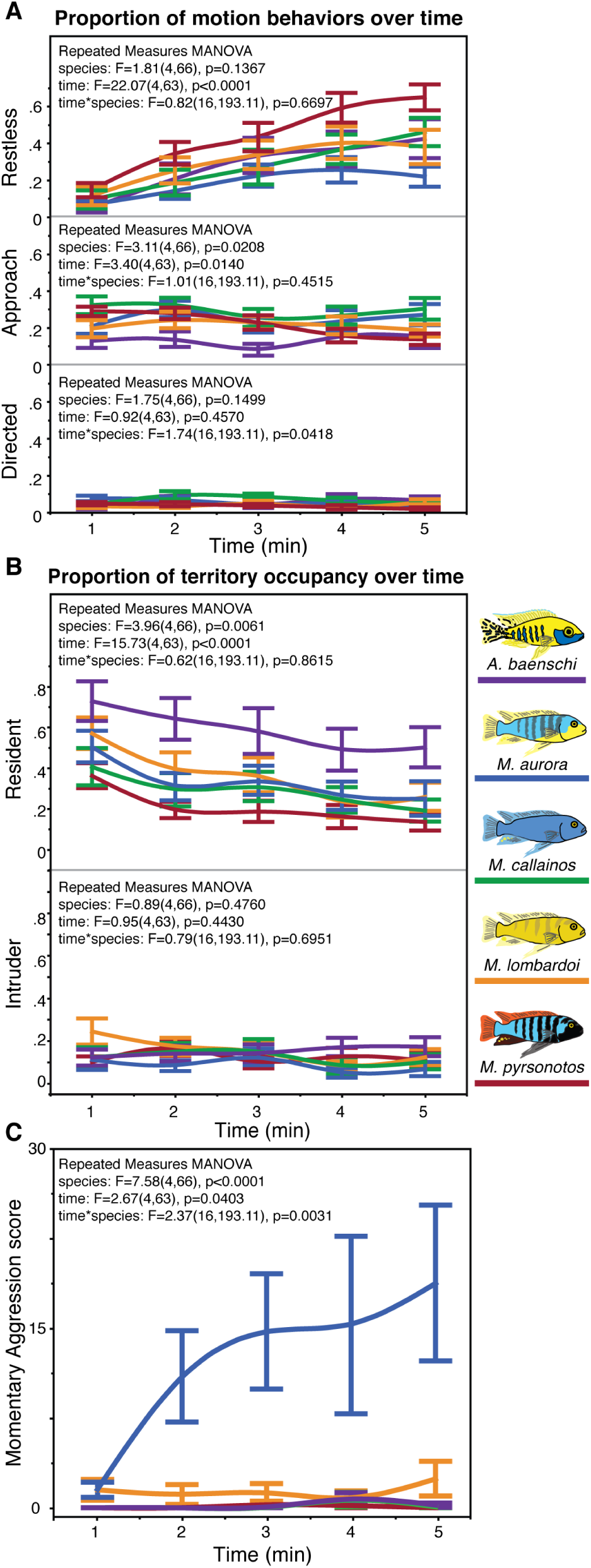
Figure S2. Behaviors over time. The proportion of time spent engaged in motion behaviors (A), and territory occupancy (B) for each minute increment over the five-minute assay. Note that some behaviors vary over time, some vary by species, and some vary by both species and time, as indicated by the repeated measures ANOVA for each panel. Momentary aggression also varies over time (C), with *M. aurora* standing out as the most aggressive. Colors indicate species, per legend.

### Resident-Intruder Assay

A digital video camera was placed in front aspect of the aquarium prior to filming, oriented to provide full view of the territory opening. From a family of size and age-matched individuals, a single *Labeotropheus trewavasae* was gently netted from his home tank, and placed in the front, open area of the resident (focal) fish’s aquarium to act as an intruder. This species was chosen because it differs in male nuptial coloration and craniofacial morphology from all five of the chosen focal species, as to avoid potential conspecific signals. Behavioral activity was video recorded for five minutes, after which the *L. trewavasae* was removed and returned to his home aquarium. The same family of *L. trewavasae* was used as intruder fish throughout the course of the experiment. Each round of testing was at least one week apart, allowing for the intruders to re-acclimate to their home tank following the assay. All behavioral tests were completed between the hours of 10:00-16:00. While there are alternatives to using a live fish as an intruder for such experiments, mirror tests necessarily do not allow for the testing of heterospecifics and data suggest that video playback is an ineffective stimulus for social response in Malawi cichlids^38^. Fish received levels of aggression comparable to that which occurs during standard co-culture housing in the lab, and were periodically monitored for stress during the assay.

Videos were analyzed for relative position to territory (resident and intruder), non-attentive movement states (restless free swim and directed free swim), attentive movement states (still and approach), aggressive movement state (chase), attentive action (fin spread display), and aggressive actions (quiver display and lunge) (Figure 5B, ethogram adapted from^39^) using the Observational Data Recorder (ODrec v2.0 beta).

### Cortisol Collection

To approximate circulating cortisol levels, we used a non-lethal method for collecting the hormone from holding water as follows (modified from (Wong et al. 2008)). After the final behavioral assay, each fish was placed in a blinded glass 600 mL beaker with 300 mL of aquaculture system water for one hour. From the original 300 mL, 200 mL of collected water was run through a single-use C-18 chromatography column (Waters) pre-flushed with 4 mL ethanol and 4 mL reverse osmosis filtered, deionized water. The extracted hormones were eluted into 4 mL 100% molecular-grade ethanol, then pelleted using a SpeedVac vacuum centrifuge. For each cohort, a control sample of system water was placed in a beaker without a fish, and processed in the same manner to determine baseline levels of hormone in the water. Cortisol was quantified using a colorimetric enzyme-linked immunoassay following manufacturer’s instructions (EIA, Cayman chemicals).

### Statistics

Basic summary values were generated in Excel. All statistical analyses were performed using packages available in JMP v12 (SAS). For examination of behaviors over time, average values for each behavioral measure were calculated in 1-minute increments, and a repeated measures MANOVA was used to model the effects of species, time, and species by time interaction. Mean summary values for eight movement and action behaviors (count or duration, where appropriate) were compared for species, sex, and interaction effects, with model p-values corrected for multiple testing using the Holm-Bonferroni method^40^. In order to evaluate levels of aggression over time, a pseudo-multivariate metric for momentary aggression was created using weighted values for each time point. Aggressive acts were weighted twice as much as attentive acts, and all acts were weighted twice as much if the fish was outside of its territory (offensive vs defensive). The proportion of offensive fin spread displays was determined by dividing the time (in seconds) that fin spread displays took place outside the territory by the total time fin spread displays were performed.

Transition probabilities were analyzed as follows: Each individual had a transition probability calculated for each behavior (for example, the ‘approach’ motion state had four other possible motion states into which it could transition, and the probability of each of those state changes was based on the total number of state changes from ‘approach’.) Those individual transitions were classified based on the type of state change (aggressive to attentive, aggressive to non-aggressive, and so on), and then averaged to determine the mean proportion of transitions types. These absolute proportions do not add up to 1, given that any individual might be missing specific transitions (for example, a fish may have been very likely to chase after approaching, but have no instances of chase after being still, both of which are types of shifts from aggressive to attentive acts).

Excreted cortisol level differences by species were analyzed using the non-parametric Wilcoxon Rank-Sum test, which is preferable given the unequal variance between species. Relationships between cortisol levels and behaviors were each modeled using a two variable linear model, and corrected for multiple testing, again, using the Holm-Bonferroni method^40^.

**Table 1.**
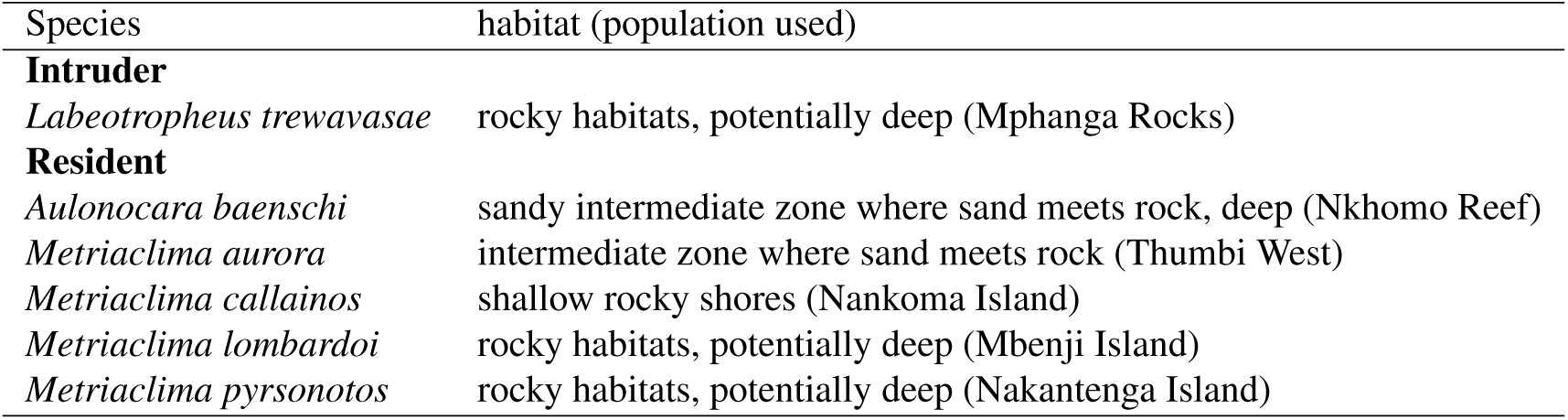
Species used in this study

## Acknowledgements

Thank you to members of the Roberts lab for animal husbandry, and to the W.M. Keck Center for Behavioral Biology at North Carolina State University for funding for this work.

## Author contributions statement

E.C.M. and R.B.R. conceived the experiment(s), E.C.M conducted the experiment(s), E.C.M analyzed the results. Both authors reviewed the manuscript.

## Additional information

The authors report no **competing financial interests**. All animal experiments were non-invasive, and conducted under the supervision of the North Carolina State University Institutional Animal Care and Use Committee (IACUC), protocol number 14-138-O.

